# Genetic and epigenetic characteristics associated with the rapid radiation of *Aquilegia* species

**DOI:** 10.1101/782821

**Authors:** Zhen-Hui Wang, Tianyuan Lu, Ming-Rui Li, Ning Ding, Li-Zhen Lan, Xiang Gao, Ai-Sheng Xiong, Jian Zhang, Lin-Feng Li

## Abstract

Elucidating the genetic and epigenetic bases underlying species diversification is crucial to understanding the evolution and persistence of biodiversity. As a well-known horticultural plant grown worldwide, the genus Aquilegia (columbine) is also a model system in adaptive radiation research. In this study, we surveyed the genomes and DNA methylomes of ten representative Aquilegia species from the Asian, European and North American lineages. Our inferences of the phylogenies and population structure revealed clearly high genetic and DNA methylomic divergence across the three lineages. By multi-levelled genome-wide scanning, we identified candidate genes exhibiting lineage-specific genetic or epigenetic variation patterns that are signatures of inter-specific divergence. We demonstrated that these species diversification-associated genetic variations and epigenetic variabilities were partially independent but were both functionally related to various biological processes vital to adaptation, including stress tolerance, cell reproduction and DNA repair. Our study provides an exploratory overview of how the established genetic and epigenetic signatures are associated with the rapid radiation of Aquilegia species.

## Introduction

Adaptive radiation is the rapid diversification of a single ancestral species into a vast array of common descendants that inhabit different ecological niches or use a variety of resources, but differ in phenotypic traits required to exploit diverse environments^1–4^. Disentangling the evolutionary mechanisms underpinning adaptive radiation is fundamental to understanding the evolution and persistence of biodiversity^5,6^. This has been a key focus of many studies which were investigating different animal and plant lineages that diversified through adaptive radiation, including Hawaiian silversword, Caribbean anoles, Darwin’s finches, and African cichlids^7–10^. However, it remains under-investigated as to why some lineages could diversify rapidly but their close relatives or other sympatrically distributed lineages did not. In the past decades, accumulating evidence from diverse radiation lineages suggest that both the extrinsic environmental factors (e.g., resource availability) and genetic variations can determine the rate and volume of species diversification^11^. Among the environmental triggers, ecological opportunity is considered as the primary mechanism that causes rapid adaptive radiation through acquisition of key innovations, penetration of new environments and extinction of competitors^2,12^. On the other hand, new species also arise as a result of new genetic variations being preserved which could ultimately influence the phenotypic disparity, where natural selection act on, among closely related species^13^. In the rapid speciation of the African cichlid fishes, extrinsic environmental factors (e.g., ecological specialization) and genetic mechanisms (e.g., adaptive introgression) acted together to provoke the repeated adaptive radiation in geographically isolated lakes^7,11,14,15^.

The genus *Aquilegia* L. (columbine) is a well-recognized model system to study the evolutionary mechanisms underlying adaptive radiation^16,17^. This genus includes approximately 70 recently diversified species that are widely distributed in the temperate zones of North America and Eurasia^18^. Phylogenetic and geographic inferences have illustrated two independent adaptive radiations of North American and European lineages from the ancestral Asian species^17,19^. For example, floral diversification of the North American *Aquilegia* species is highly correlated with the pollinator specialization^20–23^. In contrast, ecological adaptation and geographic isolation are considered as the major driving forces promoted rapid radiation of the European species^17,24^. In Asia, changes in pollinator and ecological habitats are both proposed to be the underpinning mechanisms that resulted in the diversification of more than 20 morphologically distinct species^25,26^. These Asian *Aquilegia* species constitute four highly divergent lineages corresponding to their geographic origins and have evolved relatively independently^25,26^. Despite this well-described evolutionary history and crucial role played by environmental factors, how genetic and epigenetic factors are involved in the rapid speciation in this genus remains poorly investigated.

In this study, the main objective is to survey the genomes and DNA methylomes of 36 accessions from ten worldwide *Aquilegia* species from the Asian, European and North American lineages. Among the Asian species, four phylogenetically distinct species (*A. japonica, A. oxysepala, A. yabeana, and A. viridiflora*) were selected according to their geographic distributions and ecological habitats. *Aquilegia japonica* and *Aquilegia. oxysepala* are sister species inhabiting alpine tundra and low altitude forest niches in northeastern China, respectively^25,27^. Our previous studies have documented that natural selection during ecological specialization together with genetic drift under geographic isolation caused the rapid evolution of reproductive isolation between these two species^25,28^. Here, we further investigated how diverse evolutionary driving forces shaped genetic and epigenetic architectures of the two species in the processes of speciation and adaptation. In addition, we also evaluated patterns of nucleotide variation and cytosine methylation in the *A. yabeana* and *A. viridiflora*. The former species shares highly similar morphological traits and ecological niches with the *A. oxysepala* but is allopathically distributed in northern China. In contrast, while the *A. viridiflora* is sympatrically distributed with *A. yabeana* and *A. oxysepala* in northern and northeastern China, it often occupies rocky and sandy ecological niches. As a supplementary, we also examined nucleotide and cytosine methylation variation patterns of the North American and European lineages. Our study will provide a genome-wide view of how the specific genomic and epigenomic variation patterns are correlated with the diversification of Aquilegia species.

## Results

### Population structure and nucleotide variation pattern

Neighbor-joining (NJ) trees were reconstructed for the 36 *Aquilegia* accessions based on 689,123 homozygous SNPs. The phylogenic analysis suggested that these accessions of the ten species formed three distinct lineages corresponding to their geographic origins (**Figure 1a** and **Figure S1**). In brief, all 22 accessions of the four East Asian species, *A. japonica, A. oxysepala, A. yabeana* and *A. viridiflora*, clustered as a monophyletic lineage, with the first two species and their hybrid forming a clade and the last two species grouping as a sister clade. In contrast, the West Asian species *A. fragrans* clustered with the geographically adjoining European species. The principal component analysis (PCA) and population structure inferences also revealed distinct genetic structure of the three phylogenetic lineages (**Figure 1b** and **c**). It should be noted that one *A. alpina* var. alba accession shared the same ancestral genetic cluster with the North American lineage, while the putative hybrid of the *A. oxysepala* and *A. japonica* possessed an admixed genetic background (**Figure 1b** and **c**).

**Figure 1.**
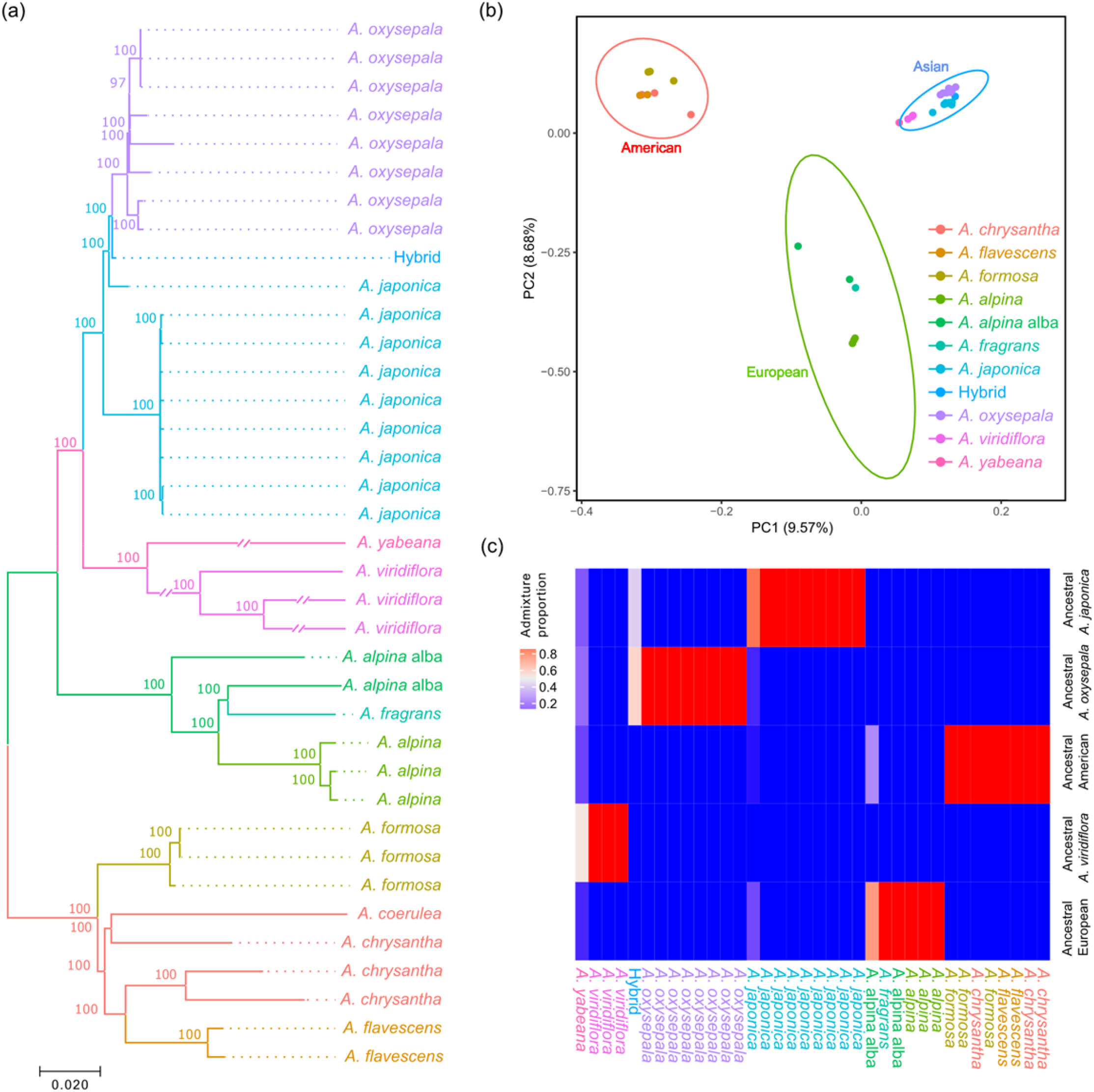
Phylogenetic relationship and population structure of the ten worldwide *Aquilegia* species. (a) Phylogenetic tree of the 36 accessions constructed by neighbor-joining algorithm based on 689,123 whole-genome SNPs. (b) PCA reveals genetic similarity within each of the three lineages and genetic disparity between lineages based on 15,988 LD-pruned SNPs. Ellipses of each lineage denote 99% confidence region estimated from distribution of the first two principal components. (c) Population admixture of the 36 *Aquilegia* accessions.

To further gain an insight into genome-wide nucleotide variation pattern of the ten *Aquilegia* species, we calculated nucleotide diversity (π) and genetic divergence (F_ST_) for each chromosome and for 100-kb sliding windows, respectively. Among the three phylogenetic lineages, the Asian *Aquilegia* species harbored the highest nucleotide diversity compared to the European and North American lineages across the seven chromosomes (**Figure S2**). By comparing the nucleotide diversity for each 100-kb sliding window, we observed a moderate correlation of genome-wide variation pattern among the three lineages (Spearman R = 0.42-0.56) and a high correlation between the *A. oxysepala* and *A. japonica* (Spearman R = 0.70) (**Figure S3**). In particular, 116 of 241 low genetic diversity genomic regions (LDGRs, with 5% lowest π) were shared by at least two of the three lineages (**Figure S4**). Between the *A. oxysepala* and *A. japonica*, while we defined 148 LDGRs and 148 high divergence genomic regions (HDGRs, with 5% highest F_ST_), only seven candidate genomic regions overlapped (**Table S1**).

### Identification of the genomic regions indicating selection pressure and highly impactful genetic variations

Candidate genes or genomic regions associated with adaptive divergence were determined from three perspectives. First, we considered genes localized within the regions that showed low intra-specific diversity but high inter-specific divergence to be representative of intra-specific genetic differences. We thus identified 23 genes from the above seven candidate genomic regions that were both HDGRs and LDGRs shared by *A. oxysepala* and *A. japonica* (**Table S1**). Genes within these genomic regions were functionally associated with meiotic nuclear division, adenine methyltransferase and basic cellar activities.

While the first strategy mainly relied on genome-wide scanning for 100-kb non-overlapping sliding window, we also employed a functional annotation-based approach to identify highly impactful conservative clade-specific variations (CCVs) from both the within and between lineage comparisons. Our results revealed that a considerable proportion (17.9-40.5%) of the CCVs were identified in the gene body regions (**Table S2**). We then examined the potential functional impacts of genes harboring these identified CCVs. Between the *A. oxysepala* and *A. japonica*, the CCV-carrying genes were enriched in several vital biological pathways related to cell reproduction, including telomere maintenance, DNA repair, and DNA helicase activity (**Figure 2** and **Table 1**). For example, two candidate genes (*Aqcoe6G160300* and *Aqcoe7G062500*) coding for *Xklp2* (*TPX2*) were functionally correlated with spindle assembly during the mitotic process (27, 28). Among the three phylogenetic lineages, the CCVs-harboring genes were also functionally involved in the mitotic chromosome condensation, DNA ligase activity and aminopeptidase activity (**Figure 2** and **Table 1**). For instance, two CCV-containing genes (*Aqcoe2G276600* and *Aqcoe1G273400*) encoding DNA mismatch repair proteins *MutS*/*MSH* and *MutS2* (ref. 29) carried one Asian-specific-to-American frameshift variant.

**Figure 2.**
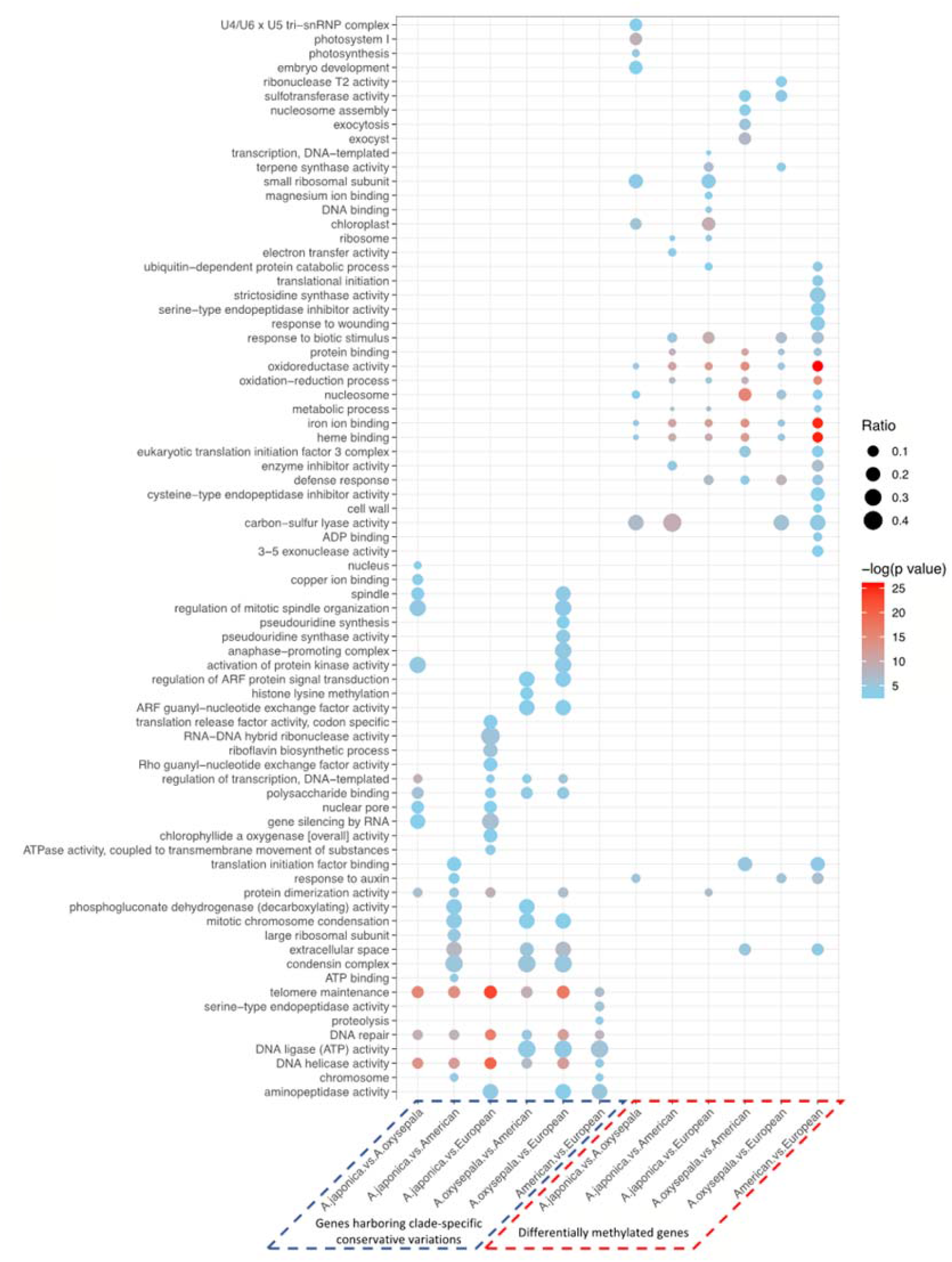
Functional enrichment of genes harboring highly impactful CCVs and DMGs. CCV-containing genes specific to either of the two lineages/species being compared were merged to construct a target gene set. Ratio denotes proportion of CCV-containing genes or DMGs in the corresponding gene set of interest. Absence of dot indicates no significant enrichment.

**Table 1.**
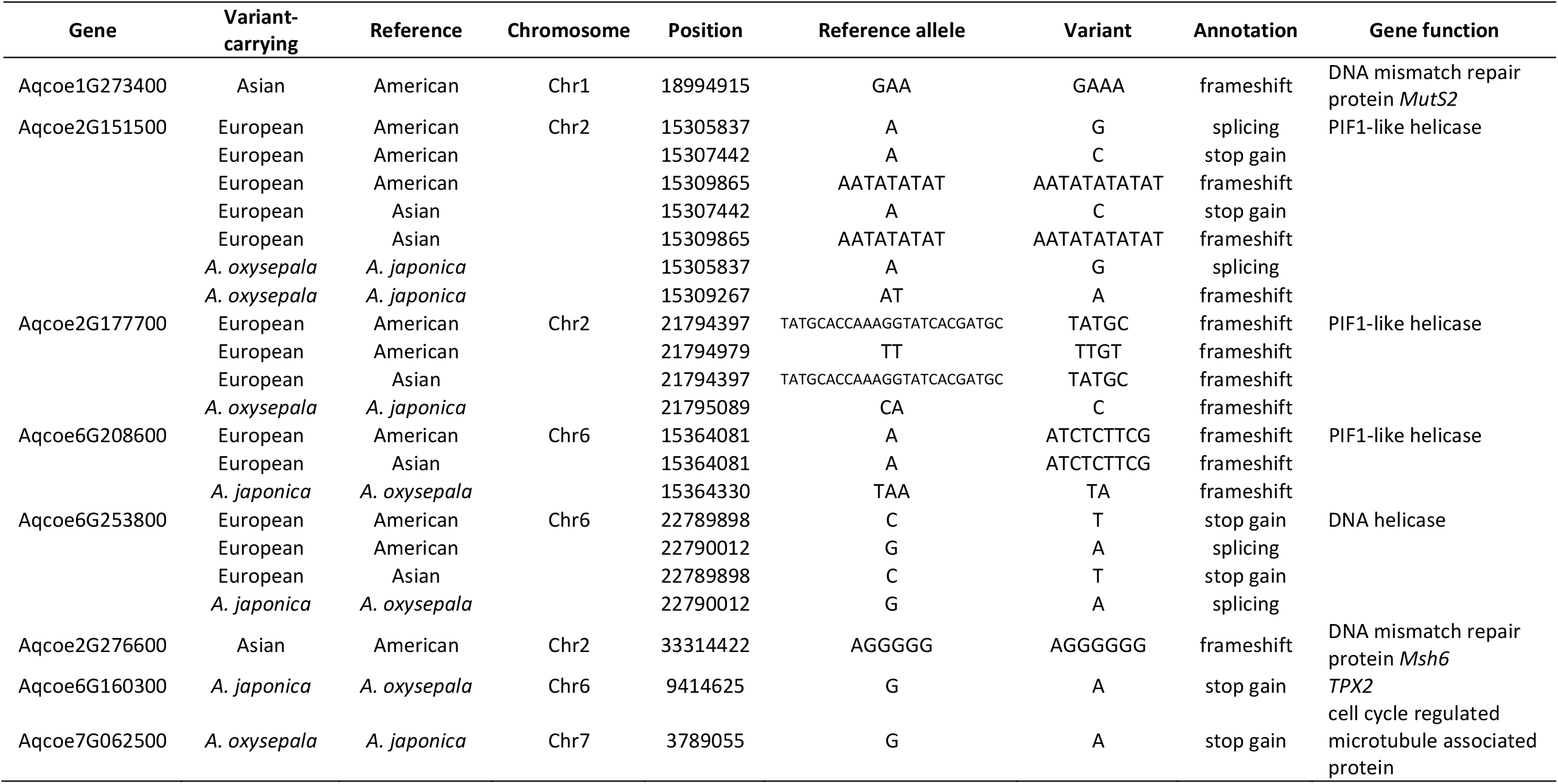
Information of the high-impact conservative clade-specific variants (CCVs) in the cell reproduction related genes.

Thirdly, we also derived pair-wise synonymous (d_S_) and non-synonymous (d_N_) mutation rate to identify genes informative of positive or purifying selection pressure. We found that species within the Asian lineage experienced significantly stronger positive (d_N_/d_S_ > 1) and purifying (d_N_/d_S_ < 1) selection pressures compared to the European and North American lineages (Wilcoxon rank sum test, all Bonferroni-corrected *p* values < 1.5×10^−16^) (**Figure S5**). Likewise, the European species showed significantly stronger purifying selection (Wilcoxon rank sum test, Bonferroni-corrected *p* value = 7.8×10^−8^) compared to the North American species.

### CG methylation patterns and differentially methylated genes

In parallel with the above genomic analyses, we also investigated CG methylation pattern of the representative *Aquilegia* species. Despite variability across the 36 *Aquilegia* accessions, the North American, Asian and European species showed no distinguishable differences (*t* test, all Bonferroni-corrected *p* values > 0.01) in overall percentage of methylated cytosines (**Figure 3a**). We then performed PCA to examine the CG-cytosine methylomic diversity of all the *Aquilegia* accessions. The resulting overall methylation pattern highly resembled the above genomic inferences, with the European and American species forming two distinct groups and the four Asian species forming three separate clusters (**Figure S6**). We then assessed the CG methylation patterns for the European and North American lineages as well as the three Asian species (*A. japonica, A. oxysepala* and *A. viridiflora*) separately. Consistent with the described genomic features, heterogeneous pattern of the CG methylation was also observed for the seven chromosomes, with the chromosome 4 demonstrating obviously higher overall CG methylation divergence compared to the other six chromosomes (**Figure 3b**). We further quantified CG methylation level deposited in the genic regions, putative *cis*-regulatory regions and CG island, respectively. In genic and regulatory regions, all three lineages shared similar modification patterns with apparent depletion of CG methylation around the transcription start site (TSS) and transcription end site (TES) (**Figure 3c**). However, the American lineage exhibited hyper-methylation (more than 10%) around the center of CG islands and a more drastic decrease throughout the CG island shores compared to the European and Asian species (**Figure 3d**).

**Figure 3.**
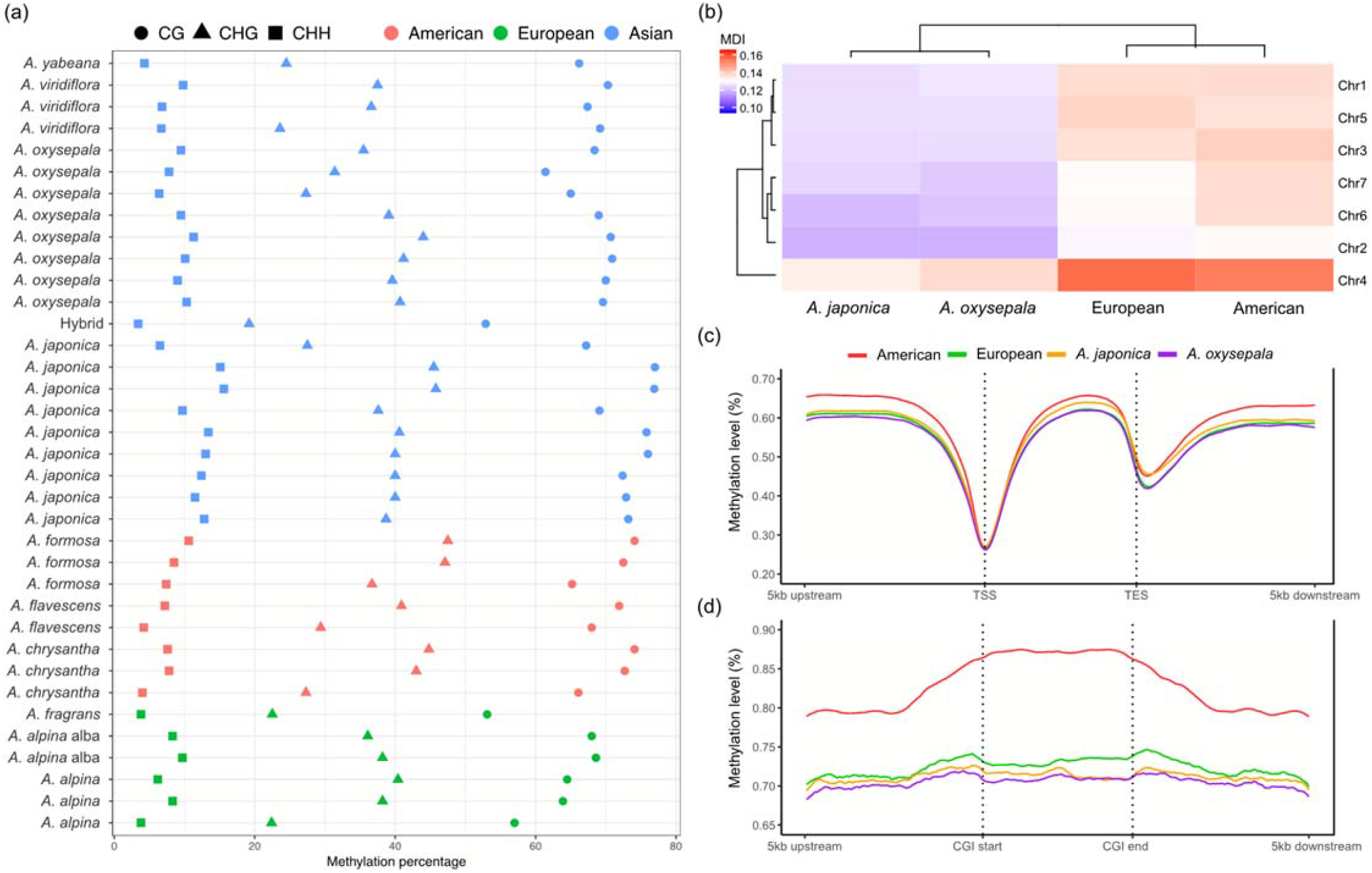
Patterns of cytosine methylation for the ten worldwide *Aquilegia* species. (a) Genome-wide cytosine methylation levels of 36 accessions. (b) MDI illustrates chromosome-level CG methylation similarity. *Aquilegia viridiflora* was used as the reference. (c) CG methylation profiling in genic region across the four *Aquilegia* groups. Each row represents one genic region starting at 5-kb upstream of its TSS and terminating at 5-kb downstream of its TES, sorted by mean methylation level of all analyzed CG loci. Gene body regions were scaled to have the same length. (d) CG methylation profiling in and around CG islands.

To examine the biological impacts of CG methylation on the species diversification, differentially methylated regions (DMRs) and differentially methylated genes (DMGs) were identified for both within- and between-lineage comparisons, respectively (**Tables S3** and **S4**). Within the Asian lineage, 3,622 DMRs in 2,899 DMGs were identified between the *A. japonica* and *A. oxysepala*. Functional enrichment of these DMGs indicated that the two species may have different activities in photosynthesis-related pathways, including photosystem I, photosynthesis and chloroplast (**Figure 2**). For example, two photosynthesis-related genes, *PsaA*/*PsaB* and *CemA*, showed significantly differential methylation between the two species in the genic regions (**Figure S7a** and **b**). At the inter-lineage level, apparently more DMGs were identified between the North American and European species (6,087 genes) compared to those of between the two lineages and Asian species (3,308-5,003 genes) (**Table S3** and **S4**). DMGs characterized from the inter-lineage comparisons were mainly involved in the plant growth (e.g., response to auxin) and defense, response to biotic stimulus and wounding (**Figure 2**).

We then examined whether the candidate genes (CCV-carrying genes and DMGs) superimposed on the same signature of natural selection. We found while a considerable proportion of the candidate genes were shared for each of the genetics- and epigenetics-based assessments (**Figure S8**), they showed a segregated distribution pattern across all comparisons (**Figure S9**). Likewise, the Gene Ontology (GO) enrichment analyses of the candidate gene identified from the genetic and epigenetic levels were enriched in functionally complementary pathways (**Figure 2**), suggesting co-existence of different underlying evolutionary mechanisms.

### Association between epigenetic variability and genetic variations

Since both genetic variation and differential CG methylation seemed to have crucial and multifaceted influences on the adaptation of the ten *Aquilegia* species, we wondered whether differential epigenetic modifications were dependent on genetic variations. Among the 588,659 CG loci examined, 224,222 (38.09%) carried a CG-loss variation. We then illustrated epigenetic variability for the variation-carrying and non-variant CG loci, respectively. As shown in Figure 4, genetic-epigenetic associations of varying magnitude were observed in both types of CG loci. The variation-carrying CG loci conveyed information that highly resembled their genetic background. The overall methylation pattern was highly conserved within the same species but exhibited obvious divergence across the ten *Aquilegia* species (**Figure 4a**). In contrast, CG methylation divergence at the non-variant CG loci varied with higher variability at both the intra- and inter-specific levels (**Figure 4b**). By examining the correlation of genetic variability and cytosine methylation, we found that CG methylation divergence at variation-carrying CG-site was largely attributable to the CG-loss variations (**Figure 4c**). In particular, 75% of the CG-loss variations occurring at the most highly variable CG-methylated dinucleotides could explain at least 75% of the total epigenetic variability *per se*. Nevertheless, there was still a considerable proportion of epigenetic variability that could not be sufficiently explained by the variant-CG locus (**Figure 4d**).

**Figure 4.**
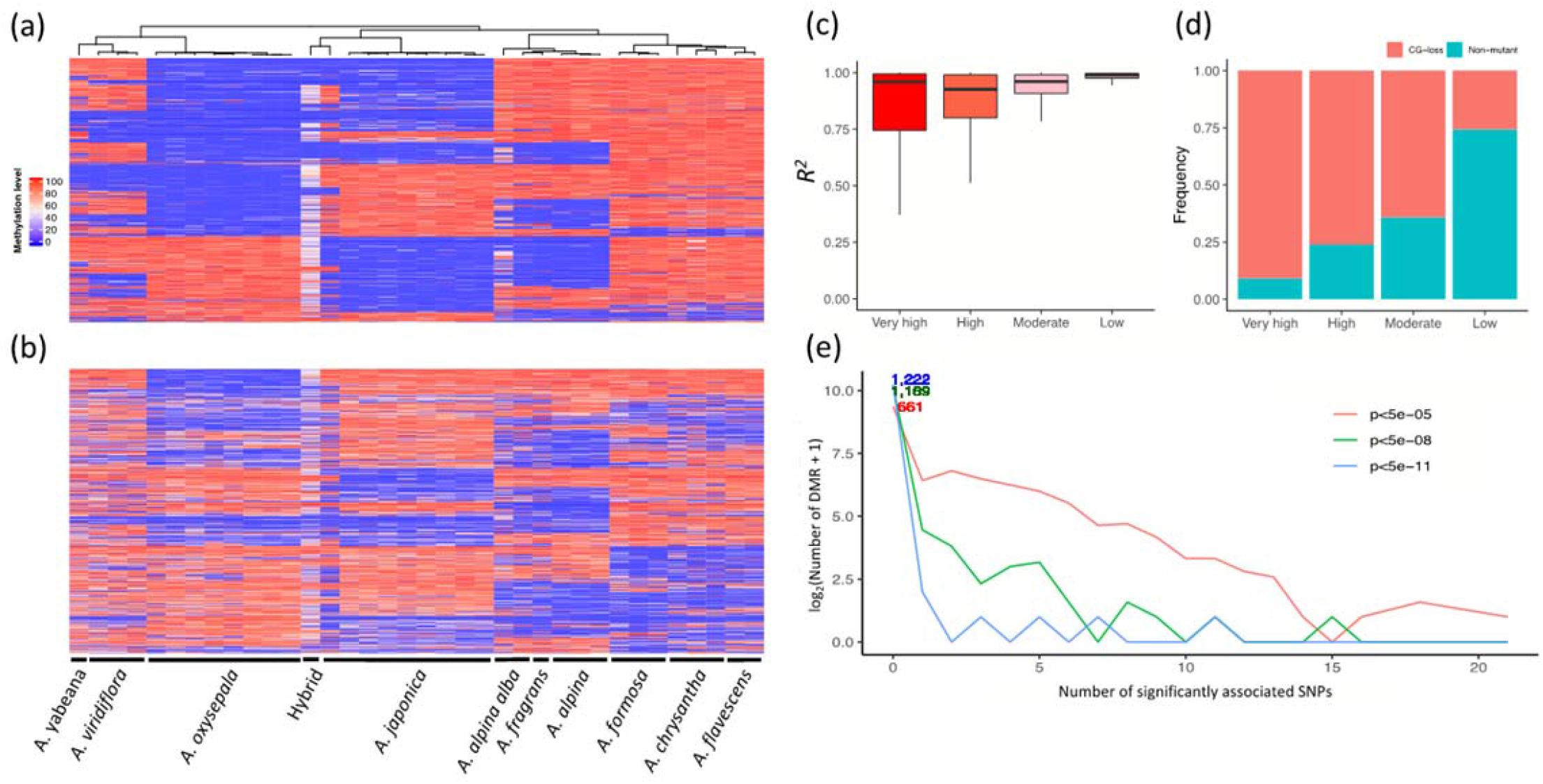
Association between CG-loss variations and epigenetic variability. (a) Top 3,000 most variable CG loci containing CG-loss variations. (b) Top 3,000 most variable non-variant CG loci across 36 accessions show clade-specific methylation patterns. CG methylation in the hybrid tends to be neutralized possibly due to heterozygosity. (c) Linear regression demonstrates that CG-loss variations explain a large proportion of CG methylation variation. (d) Summary of composition of each category with regard to whether each CG locus contains a CG-loss variation. Epigenetic variability was determined by standard deviation in methylation β value across all 36 accessions. CG loci with top 10,000, 10,001-50,000 and 50,001-150,000 largest standard deviation were ordinally labelled as possessing “very high”, “high” and “moderate” variability respectively. The rest CG loci were labelled as possessing “low” variability. (e) Association test shows most DMRs were independent of *cis*-acting SNPs. Results under different significance levels are compared in this exploratory analysis.

We also attempted to identify *cis*-driver mutations for each of the 1,229 DMRs between the *A. japonica* and *A. oxysepala*. Our results revealed that only 568 out of the 1,229 (46.2%) DMRs were significantly associated with at least one genetic variation inside or around a 500 base-pair (bp) upstream/downstream genomic region, even under the least stringent *p* value threshold (5×10^−5^), indicating that the epigenetic changes were only partially dependent on *cis*-genetic driving mutations (**Figure 4e**). Moreover, we observed weak yet significant associations between differential CG methylation and selection pressure. In most inter-lineage comparisons, DMGs were significantly more prone to be under positive selection than non-DMGs (**Table 2**), suggesting that epigenetic modifications could probably assist selection pressure in shaping genotypes. In contrast, DMGs were significantly less prone to be under purifying selection (**Table 2**).

**Table 2.** Significant correlation between differential methylation and natural selection.

| Type of selection | Differential methylation | Jap-Oxy * | Jap-Ame | Jap-Eur | Oxy-Ame | Oxy-Eur | Ame-Eur |
| --- | --- | --- | --- | --- | --- | --- | --- |
| Positive selection | DMG | 7.2% | 7.3% | 11.9% | 6.7% | 8.4% | 8.9% |
|  | non-DMG | 4.4% | 5.0% | 5.6% | 4.7% | 5.4% | 5.4% |
|  | *p* value | 0.11 | 7.3e-02 | 3.9e-05 | 6.7e-02 | 1.8e-02 | 2.8e-04 |
| Purifying selection | DMG | 3.1% | 1.8% | 2.4% | 2.3% | 2.0% | 1.9% |
|  | non-DMG | 4.3% | 4.3% | 4.7% | 4.9% | 5.1% | 4.0% |
|  | *p* value | 0.53 | 3.2e-02 | 9.1e-02 | 1.3e-02 | 8.4e-03 | 1.0e-02 |

\*: Each percentage represents the proportion of genes belonging to either DMGs or non-DMGs compared between the two corresponding clades that are under corresponding or higher strength of positive selection. For example, 7.2% indicates that 7.2% DMGs compared between *A. japonica* and *A. oxysepala* are under strong selection; 4.4% indicates that 4.4% non-DMGs compared between these two species are under strong selection. *p* values were obtained from Chi-square tests and were not adjusted for multiple testing due to dependence arising from overlapping gene sets.
Jap: *A. japonica*; Oxy: *A. oxysepala*; Ame: American; Eur: European

## Discussion

### Genetically determined mechanisms associated with the rapid diversification of *Aquilegia* species

Elucidating evolutionary mechanisms underpinning species diversification is crucial to understanding the evolution and persistence of biodiversity^2,5,6^. The genus *Aquilegia* provides an ideal system to address how the diverse evolutionary mechanisms promoted rapid adaptive radiation^16,17^. Although various environmental conditions related to ecological opportunities, such as shifts in pollinator and habitat, have been proposed to facilitate the evolution of reproductive isolation^21,25^, genetic basis associated with the rapid diversification of *Aquilegia* species has still remained largely unclear. In this study, we surveyed the genomes of ten worldwide *Aquilegia* species to address whether specific genetic architectures have been involved in the rapid species diversification. Broadly consistent with previously inferred phylogeny^17,19,25,30^, the ten *Aquilegia* species from Asia, Europe and North America formed three phylogenetically independent lineages corresponding to their geographic origins. This attribute renders the *Aquilegia* species a suitable system to identify genomic variations associated with the repeated adaptive speciation by extensive comparisons from different facets.

It has been proposed that if a genetic factor is the potential determinant promoting adaptive speciation, one would expect to identify specific genetic architectures in the diversified lineages^8,14,31^. In Darwin’s finches, for example, polyphyletic topology was observed as a general pattern in 14 morphologically distinct species, phenotypic diversity of the beak shape was mainly determined by natural selection acting on the *ALX1* gene during the ecological specialization process^8^. A similar phenomenon was observed in the East African cichlid fish where the radiating lineages are more dynamic in terms of gene content and transcriptomic landscape compared to their non-radiating relatives^14,31^. In this study, the genome-wide nucleotide variation pattern highly reflects the evolutionary history that the Asian, European and North American *Aquilegia* species have clearly diverged and evolved allopatrically in respective geographic regions. This also suggests that a considerable proportion of the genetic variations and changes in environment are intertwined during the diversification process. As expected, our genome-wide scanning for selection signatures revealed distinct positive and purifying selection modes in the intra- and inter-lineage comparisons. More importantly, the CCV-carrying genes identified from the three lineages are associated with cell reproduction (e.g., telomere maintenance and mitotic chromosome condensation) and other functionally important traits. Similar genomic feature was also observed in *A. japonica* and *A. oxysepala*. Our previous studies have demonstrated that natural selection and genetic drift together resulted in the rapid evolution of reproductive isolation^25,30^. Here, we further demonstrate that candidate genes involved in the adaptive speciation are functionally enriched in the pathways related to cell reproduction (e.g., telomere maintenance), stress tolerance (e.g., response to wounding) and basic cellular activities. It should be noted that although a majority of the enriched pathways are specific to each comparison, enrichment of cell reproduction-related pathways (e.g., telomere maintenance, DNA repair and DNA helicase activity) and stress tolerance are shared in the intra- and inter-lineage comparisons. Taken together, these findings indicate that specific genetic determinants might have conferred high adaptability to the Aquilegia species to cope with different environmental conditions.

### Evolutionary potential of cytosine methylation in the adaptation of *Aquilegia* species

The role of epigenetic modification in the long-term evolutionary process has long been debated^32–34^. It has been proposed that epigenetic variations are frequently under the genetic control which can alter rapidly as a result of environmental induction and stochastic epimutation^35,36^. Nevertheless, it has also been recognized that some epigenetic variations can persist over generations and be highly correlated with phenotypic diversity^32^. As illustrated in Arabidopsis, changes in cytosine methylation can produce meiotically stable epialleles, which could eventually lead to phenotypic diversity in the absence of genetic variations^37–39^. Here, we assessed whether the epigenetic modifications were also associated with the adaptive speciation of the *Aquilegia* species. Consistent with the genomic features detailed above, high divergence of cytosine methylation was observed across the Asian, European and North American lineages. Notably, differential cytosine methylation was not only found across the seven chromosomes but also evident in the gene body of DMGs and CG island region among the three lineages. Particularly, functional enrichment analyses identified significant associations with adaptation-related traits, including plant growth, stress tolerance and basic cellular activities. For example, the candidate DMGs identified between the *A. japonica* and *A. oxysepala*, showed significant enrichment in pathways related to diverse important phenotypic traits, such as photosynthesis, embryo development and response to auxin. These features indicated that epigenetic factors might also play a role in response to diverse environmental conditions.

We noted that some candidate genes and enriched pathways had shared hotspots of both genetic and epigenetic disparities, especially those related to cell reproduction, plant growth and stress tolerance. Many studies based on human and mouse have shown that genetic variations can manipulate *cis*-CG methylation at specific loci to further influence phenotypes, where CG methylation serves as a mediator^40,41^. By analyzing the associations between genetic and epigenetic variability, we conclude that while many CG-loss variations can directly lead to depletion of CG methylation, a lot of DMRs are not manipulated by any *cis*-variations. Since gene body CG methylation in plants generally stabilizes gene expression and is positively correlated with gene expression^42–45^, differential methylation in our study is indicative of likely differential amount of gene products. Based on these attributes together with the plausible associations between differential methylation (e.g., DMGs) and positive selection (e.g., d_N_/d_S_), we propose that epigenetic modification may be a complementary mechanism facilitating phenotypic diversity of the *Aquilegia* species.

### Limitations and future directions

We realized that this study has some limitations. Fistful, the small sample size in this study may introduce bias and inflation of false positives, and we postulate that our findings should be interpreted carefully and considered exploratory. When the association between genetic divergence and evolutionary events is investigated, it is impossible to deny the roles of other evolutionary forces. We acknowledge that the lineage-specific allele frequencies are possibly consequences of genetic drift, and genetic hitchhiking may lead to identification of candidate genes residing in neighboring genomic regions representing the other driving forces. Therefore, we claim that the candidate genes identified to be associated with adaptive radiation do not necessarily point towards causal evolutionary mechanisms. They may also be by-products of the long-term process of adaptive radiation. In addition, we never than less only focused on analysis of CG methylation as puzzles persist regarding the functional roles of CHG and CHH methylation. We also expect that future studies with larger sample sizes will be able to improve the statistical power and investigate *trans*-genetic control.

## Methods and Materials

### Sample collection, DNA extraction and whole-genome sequencing

In this study, a total of 36 accessions from ten worldwide *Aquilegia* species were collected (**Table S5**). Among the Asian species, four phylogenetically distinct species (*A. japonica, A. oxysepala, A. yabeana*, and *A. viridiflora*) are selected according to their geographic distributions and ecological habitats. *Aquilegia japonica* and *A. oxysepala* are sister species inhabiting alpine tundra and low altitude forest niches in northeastern China, respectively^25,46^. Eighteen accessions were collected to represent these two Asian species and their putative hybrid. In addition, four accessions were collected from the other two Asian species, *A. yabeana* and *A. viridiflora*. The former species shares highly similar morphological traits and ecological niches with *A. oxysepala* but is allopatrically distributed in the northern China. In contrast, *A. viridiflora* is sympatrically distributed with *A. yabeana* and *A. oxysepala* in the northern and northeastern China, but often occupies rocky and sandy ecological niches. Furthermore, six and eight accessions were sampled from the European and North American lineages, respectively. All the 36 accessions were grown in green house under the same conditions (25°C/12 hours, 16°C/12 hours). Genomic DNA was extracted from fresh mature leaves using TianGen plant genomic DNA kit. Whole genome resequencing and bisulfite sequencing were performed on the extracted genomic DNA using the Illumina X-ten platform (Illumina, California, USA). Short-insert (350 bp) DNA libraries of all accessions were constructed by NovoGene (NovoGene, Tianjin, China). Genome assembly of an admixed species *A. coerulea* “Goldsmith” was obtained from Phytozome v12.1 (https://phytozome.jgi.doe.gov) as the reference genome^16^.

### Sequence assembly, functional annotation and genetic diversity

Whole genome sequences of each accession were aligned against the reference genome using default settings of the BWA-MEM algorithm implemented in Burrows-Wheeler Aligner (BWA)^47^. Raw assemblies were realigned using IndelRealigner provided in the Genome Analysis Tool Kit by default settings^48^. Single nucleotide polymorphisms (SNPs) and insertions/deletions (INDELs) were reported using SAMtools^49^. Only the high-quality variants (SNPs and INDELs) (read depth > 3, mapping quality > 20 and missing allele < 1%) were retained for subsequent population genomics analyses. Genomic annotation of the identified variants was reported for each of the 36 samples separately. Functional annotation of each identified variant was performed using SnpEff, based on the reference genome^50^.

To infer the phylogenetic relationship between the ten *Aquilegia* species, NJ trees were reconstructed for each chromosome and the whole genome dataset using MEGA 7(ref. 51). PCA was carried out to examine the genetic diversity of the 36 *Aquilegia* accessions^52^. Ancestral components were estimated using ADMIXTURE^53^ with different number of populations ranging from one to ten. Optimal population composition with the least 5-fold cross-validation error was selected to decompose ancestral admixture. To obtain the genome-wide nucleotide variation pattern, nucleotide diversity (π) and genetic differentiation (Weir and Cockerham’s F_ST_) were calculated for each 100 kb non-overlapping sliding window using VCFtools^54,55^. Pair-wise non-synonymous-to-synonymous (d_N_/d_S_) ratios of the ten species were inferred by yn00 program in the Phylogenetic Analysis by Maximum Likelihood (PAML) package^56^. Inter-lineage d_N_/d_S_ value for each gene was derived by averaging d_N_/d_S_ values obtained from all pairwise comparisons of samples belonging to the two lineages under investigation. Candidate genes with the 5% highest and 5% lowest d_N_/d_S_ values were considered to have undergone strong positive and purifying selection, respectively.

### Cytosine methylation pattern and epigenetic population structure

Whole genome bisulfite sequencing data were pre-processed using TrimGalore (https://www.bioinformatics.babraham.ac.uk/projects/trim_galore/, accessed August 21, 2018). Paired-end reads were then aligned to the reference genome using Bismark^57^ with a moderately stringent minimum-score function (L,0,-0.3). De-duplicated alignments of the 36 *Aquilegia* accessions were used to report cytosine methylation level using “Bismark_Methylation_Extractor”, on loci with a read depth ≥ 3. Genomic annotations of the methylated cytosine site were identified based on the reference genome using an in-house Python script. PCA was conducted for 588,659 loci which were passed the quality control to infer the CG-methylomic diversity of the ten *Aquilegia* species. Differential cytosine methylation was determined at the gene and chromosome levels, respectively. At the gene level, we determined DMRs for each 100 bp non-overlapping sliding window using Cochran-Mantel-Haenszel (CMH) test to account for imbalanced read depth (Supplementary Notes). Genomic regions that possessed a Benjamin-Hochberg adjusted *p* value < 0.05 and showed inter-specific or inter-lineage methylation divergence higher than 25% were defined as significant DMRs. Genes with > 20% of the genic region being DMR(s) were defined as DMGs. Chromosome level methylation patterns were measured by chromosomal methylation discrepancy index (MDI)^58^. Methylation patterns of the identified DMGs were visually confirmed on Integrative Genomics Viewer^59^ prior to downstream analyses and biological interpretation. In addition, we identified CG islands from the *A. coerulea* “Goldsmith” reference genome using EMBOSS cpgplot with default settings^60^. Only the identified CG-enriched genomic regions with > 200 bp were defined as CG islands. We then investigated inter-specific and inter-lineage methylation patterns in and around the CG islands.

### Associations between the genetic variation and cytosine methylation

We tested for associations between the identified DMGs and genes under positive selection by a Chi-square test. Linear regression model was adopted to measure the direct causal effect of CG-loss variation on CG methylation. To further assess whether genetic variations drive the establishment of DMG, driving mutations of DMRs between the *A. japonica* and *A. oxysepala* were identified using an Eigenstrat-based method (See Supplementary Notes for more details)^61^.

### Identification of conservative clade-specific variant

CCVs were defined as variants that had a SnpEff-predicted “high” functional impact and that were conserved across all samples belonging to the same species or lineage, but not present in any sample of the other species/lineages. Since the biological consequences of heterozygous variants were less affirmable, only the homozygous point mutations and INDELs were included in the characterization of CCVs, including frameshift, stop-gain, stop-loss, start-loss and splicing-alteration variations.

### Functional analysis

The above mentioned genetic and epigenetic analyses helped to identify relevant candidate genes, which might be associated with the rapid diversification of the *Aquilegia* species from different perspectives. These candidate genes were employed to conduct functional enrichment analyses using the R package topGO with default settings^62^. Enriched GO terms that possessed a *p* value <0.05 were considered statistically significant. Since the statistical tests performed by topGO are not independent, multiple testing correction does not apply here^62^. Structures of functional domains of targeted genes were determined based on the InterPro database (https://www.ebi.ac.uk/interpro, accessed January 25, 2019). Distribution patterns of the identified candidate genes and their related functional pathways were visualized using the R package jvenn^63^.

## Data availability

All data generated from the study were submitted to EBI under the accession number PRJEB34182. All scripts for conducting computational analyses are available upon reasonable request to the corresponding author.

## Author Contributions

L.F.L., J.Z and A.S.X conceived this project. Z.H.W, T.L. and M.R.L. developed statistical analysis pipeline. Z.H.W., T.L., M.R.L., N.D., L.Z.L., Y.J.H and X.G. carried out experiments and analyzed the data. T.L., M.R.L., N.D., Z.H.W., L.Z.L., X.G., and L.F.L. participated in discussion and interpreted the data. Z.H.W, T.L. and L.F.L. wrote the manuscript. All authors read and approved the manuscript.

## Acknowledgments

We are very grateful to Aköz Gökçe for her constructive comments on our manuscript. We appreciate Dr. Peng Jiang, Zhang Zhang, and the USDA for kindly providing the seeds of the *Aquilegia* species.

## Supplementary files

**Figure S1.**
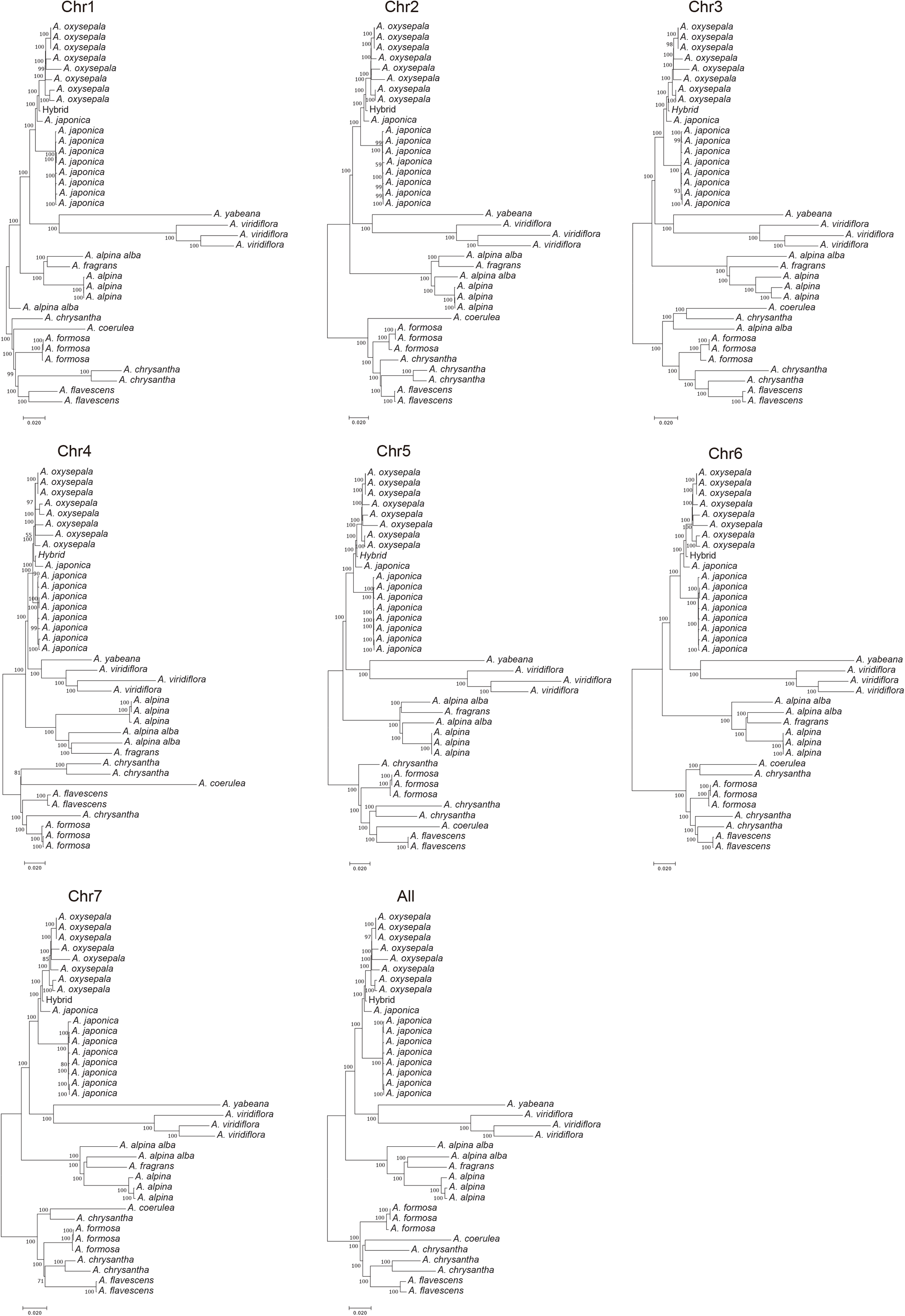
Per-chromosome phylogenetic trees reconstructed using neighbor-joining algorithm. Polymorphisms detected on each chromosome were retrieved separately to infer the phylogeny.

**Figure S2.**
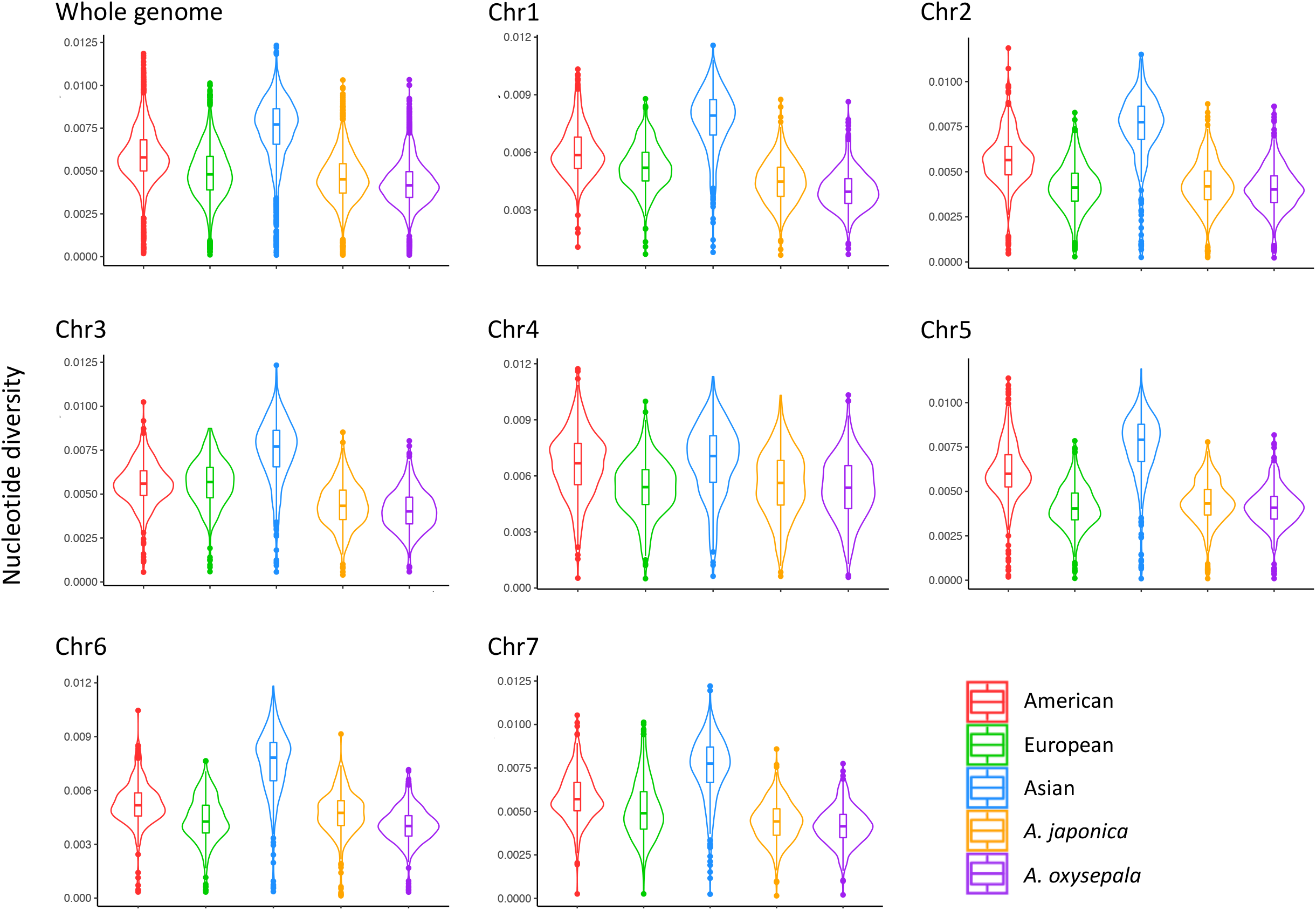
Distribution of nucleotide diversity (π) at the whole-genome level and the per-chromosome level. Nucleotide diversity was estimated for each lineage pooling corresponding species, as well as for *A. japonica* and *A. oxysepala*.

**Figure S3.**
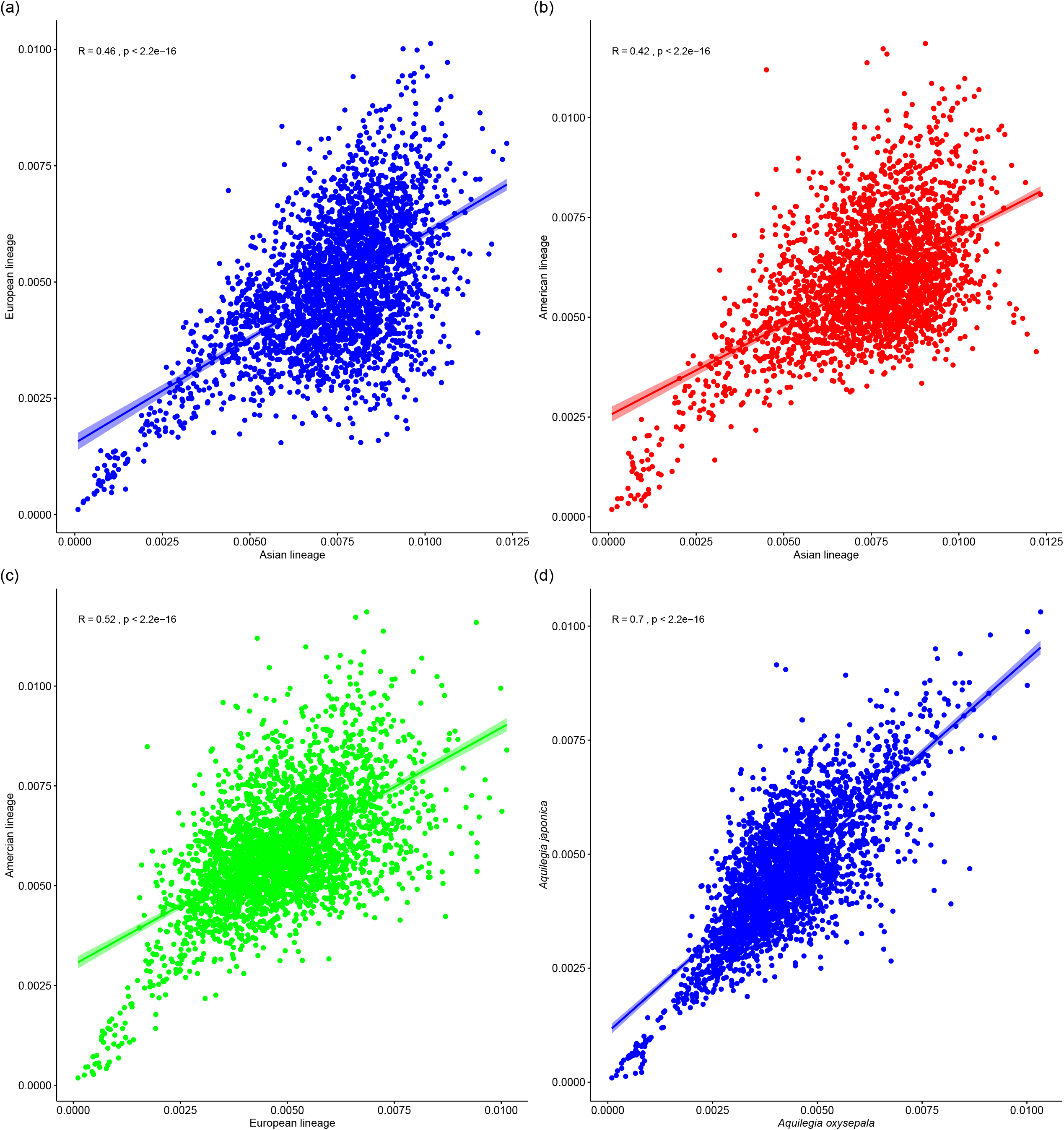
Spearman correlation of the genome-wide nucleotide variation pattern for each 100-kb sliding window between European and Asian (a), North American and Asian (b), European and North American (d), *Aquilegia japonica* and *A. oxysepala* (d). Each dot represents a 100-kb sliding window. Values on the x- and y-axis are the nucleotide diversity (π) for each sliding window.

**Figure S4.**
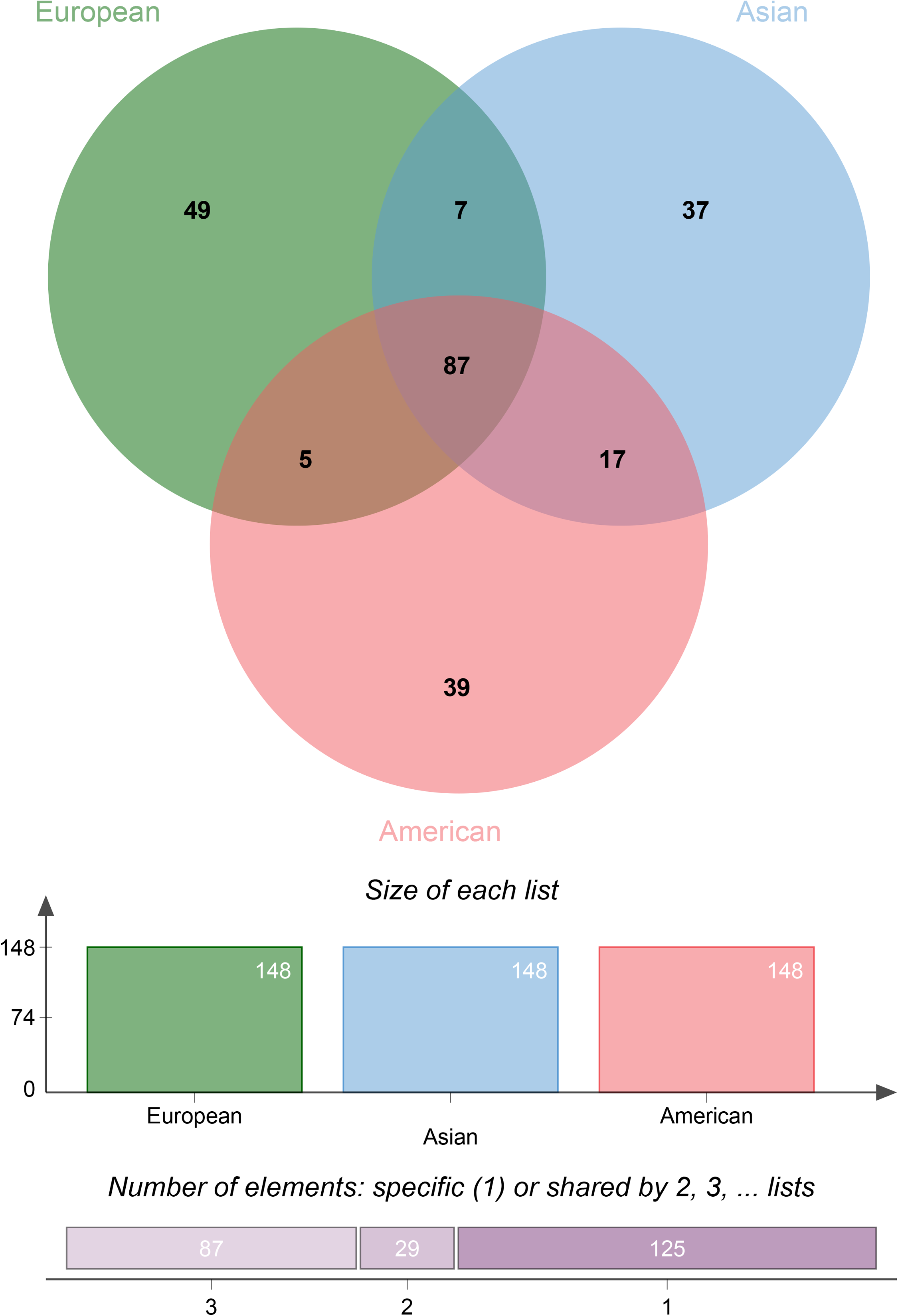
Overlapping of low diversity genomic region (LDGR) between the three lineages. 148 LDGRs with 5% lowest nucleotide diversity were defined as LDNRs in each lineage, totaling 241 unique regions.

**Figure S5.**
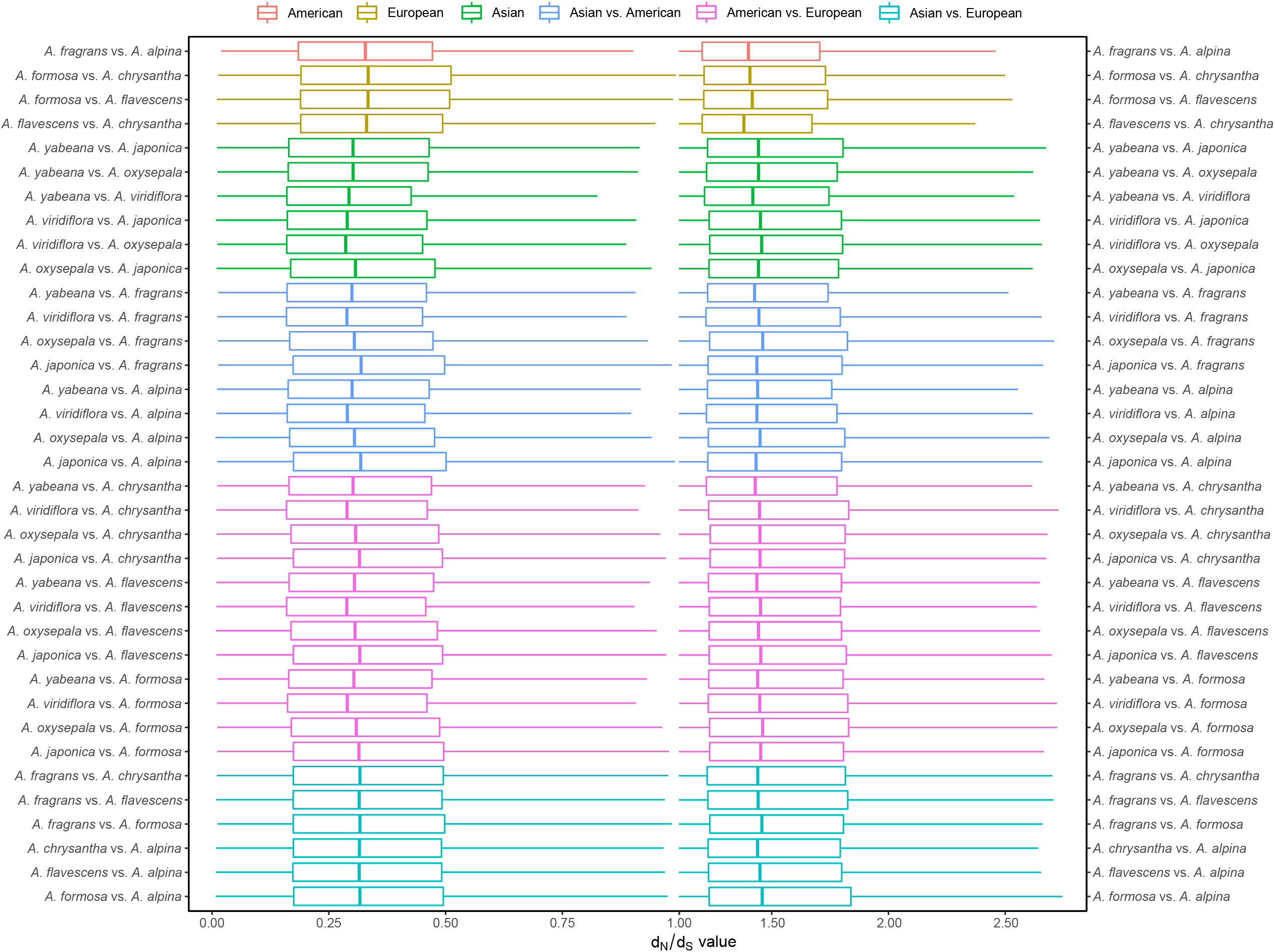
Pair-wise d_N_/d_S_ ratio for all genes between the species within and between the Asian, European and North American lineages.

**Figure S6.**
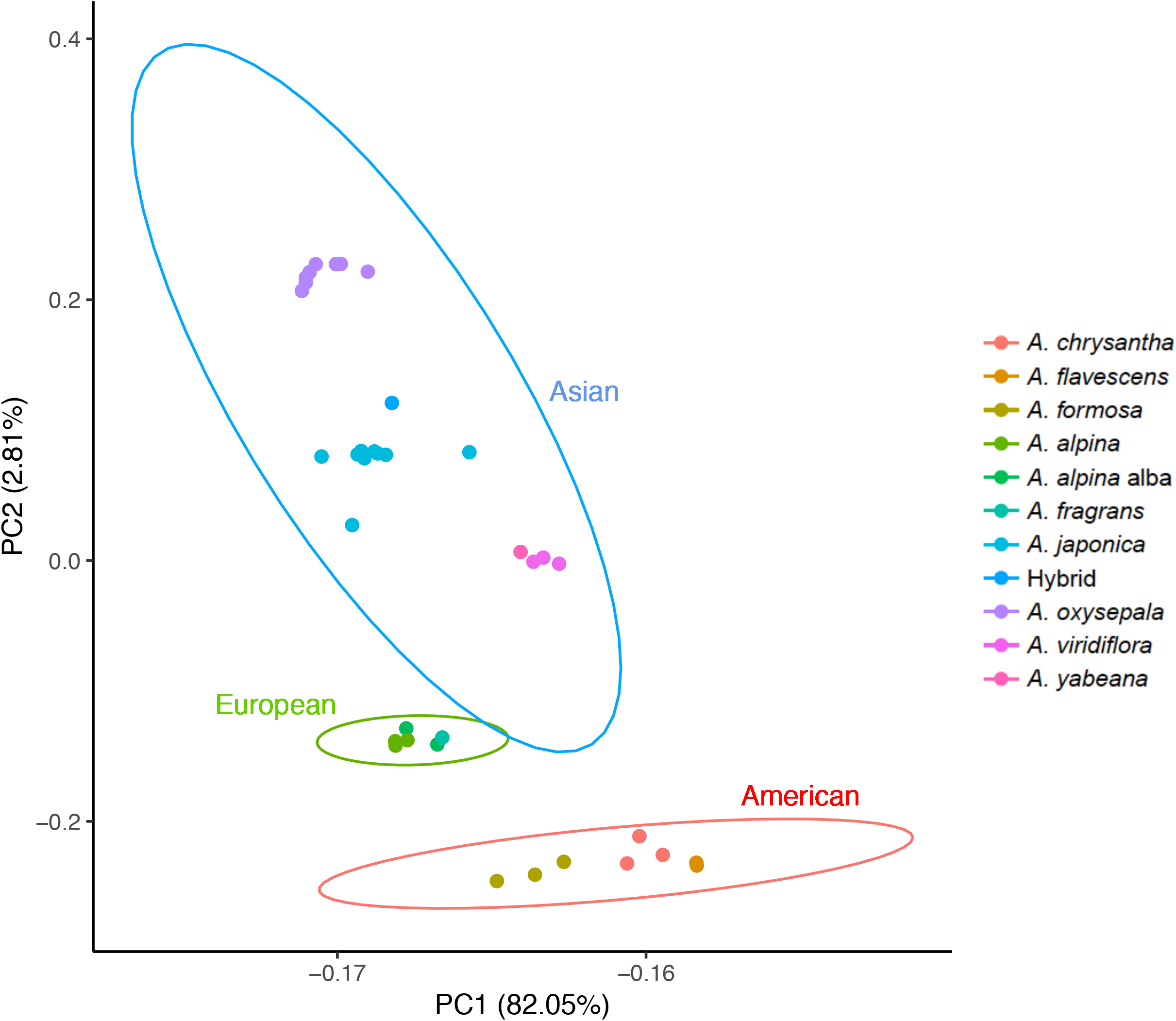
PCA illustrates three distinct clusters corresponding to the three lineages. Asian species further demonstrated higher inter-specific divergence than the American and the European species. PCA was performed based on 588,659 loci with sufficiently high sequencing quality.

**Figure S7.**
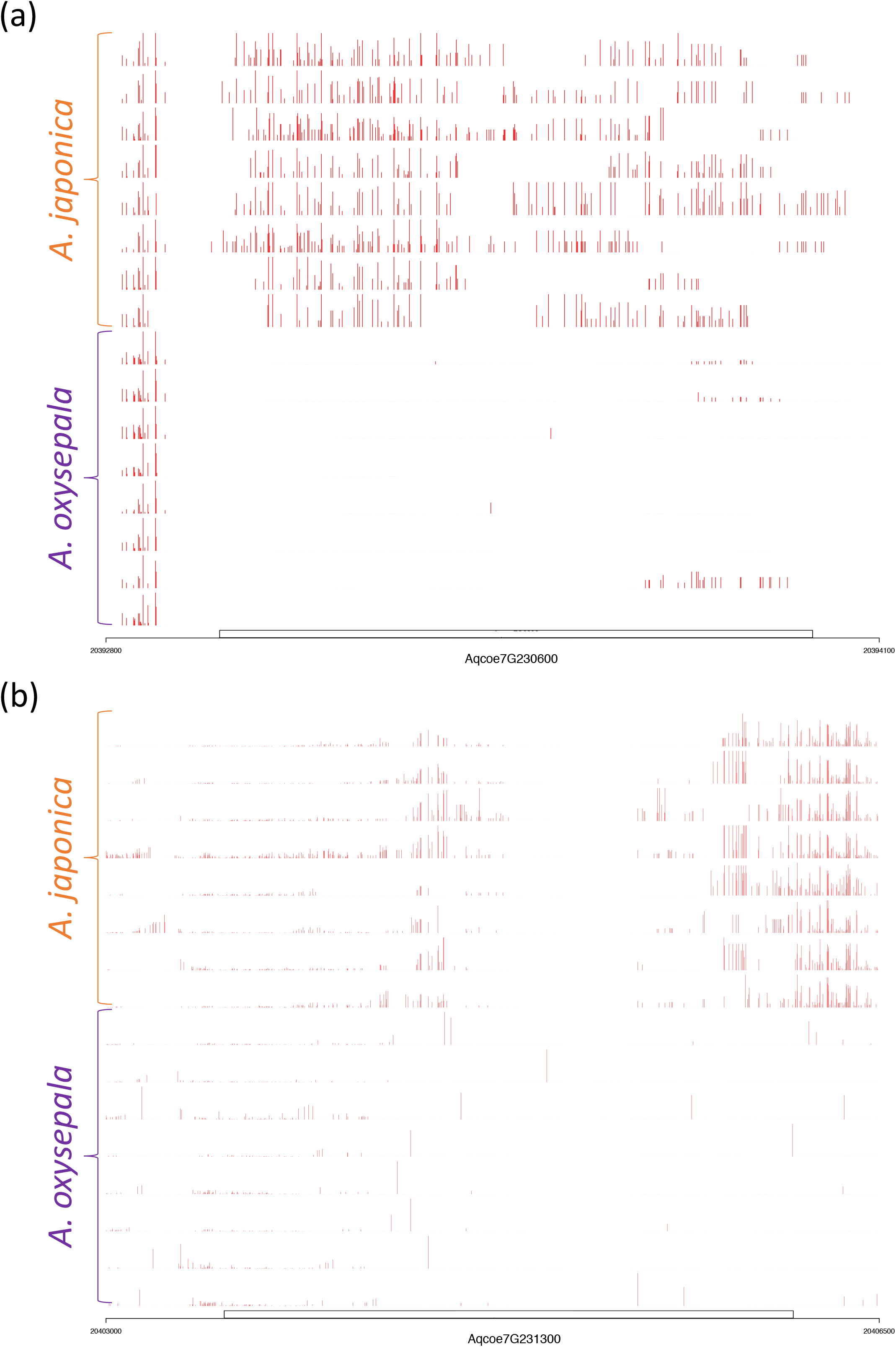
Illustration of differential methylation in two photosynthesis genes. CG methylation pattern of two genes, *Aqcoe7G230600* photosystem I *PsaA*/*PsaB* (a) and *Aqcoe7G231300 CemA* (b) in *A. japonica* and *A. oxysepala* throughout the gene body region. Red bars indicate methylation level (0-100) at CG loci. Genomic coordinates on the chromosome 7 are annotated.

**Figure S8.**
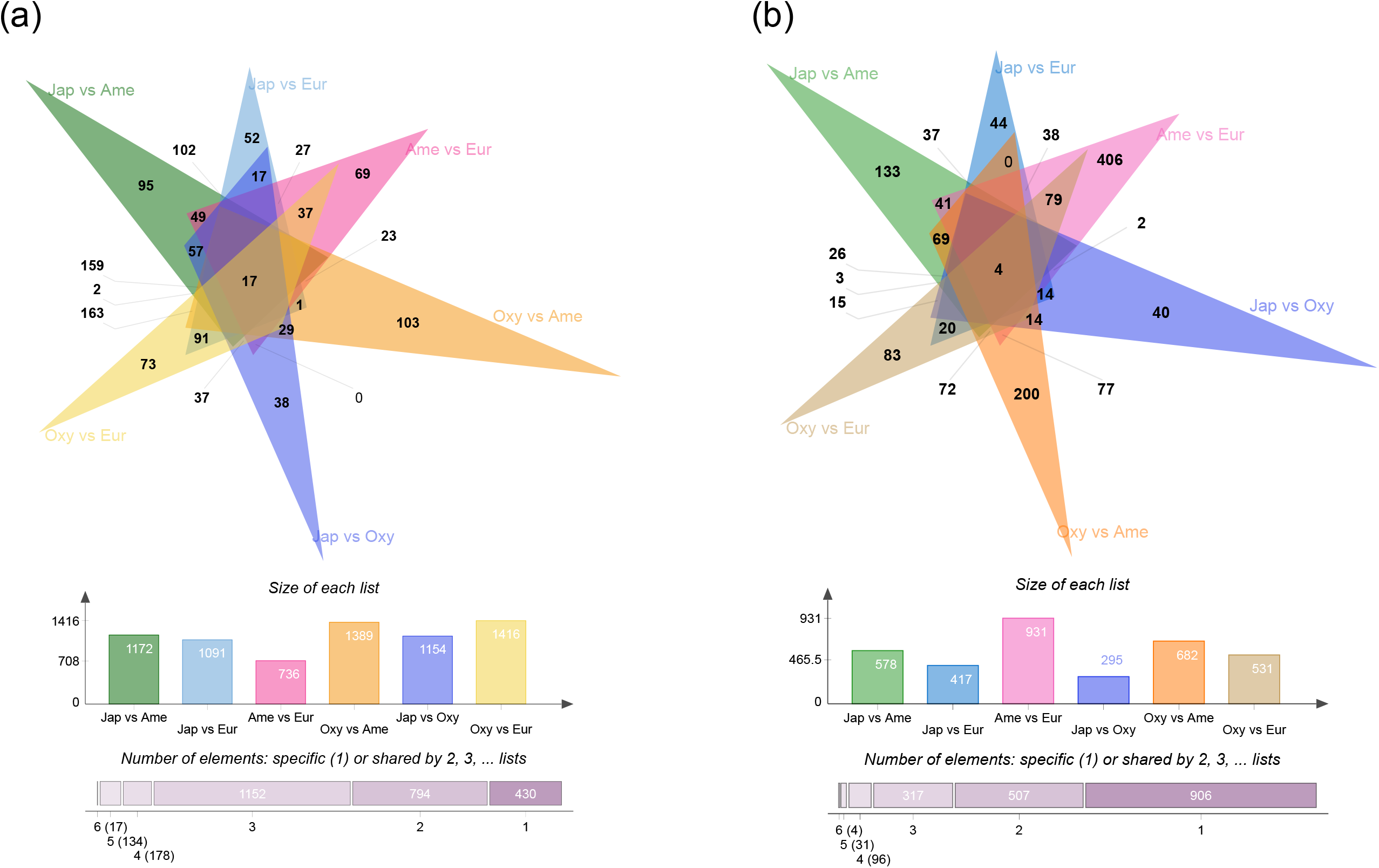
Overlapping of the CCVs (a) and DMGs (b) identified in inter-lineage/species comparisons. A considerable proportion of these CCVs (84.1%) and DMGs (51.3%) were shared by two or more inter-lineage/species comparisons.

**Figure S9.**
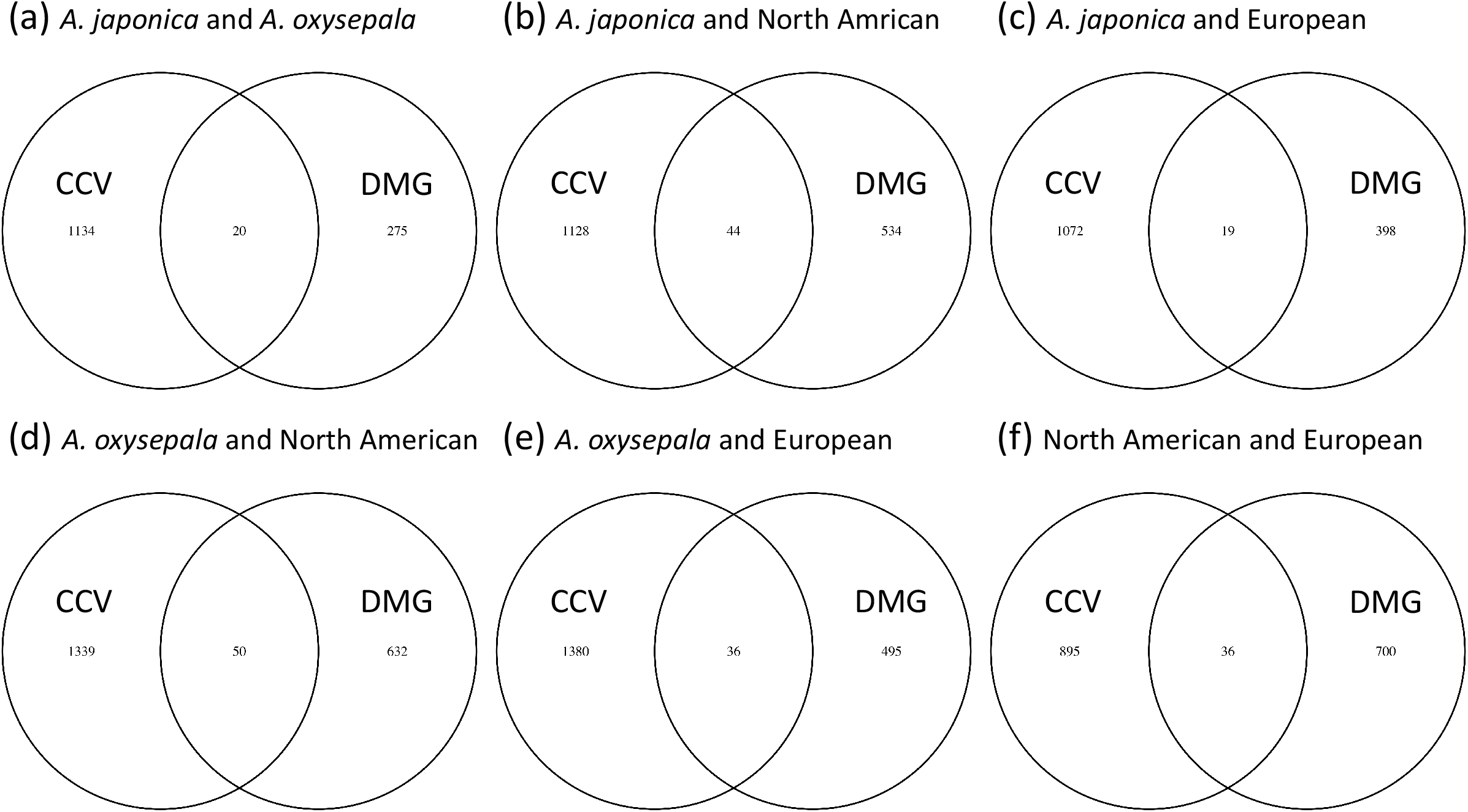
Venn analyses of the candidate genes carrying CCVs and DMGs. Each subpanel indicates the comparison between the *A. japonica* and *A. oxysepala* (a), *A. japonica* and North American (b), *A. japonica* and European (c), *A. oxysepala* and North American (d), *A. oxysepala* and European (e), North American and European (f).

**Table S1**. Candidate genomic regions that showed high genetic divergence (top 5% highest F_ST_) between *Aquilegia japonica* and *A. oxysepala* and low nucleotide diversity (top 5% lowest π) within each species.

**Table S2**. Summary of the highly impactful clade specific variations (CCVs) at both the species and lineage levels.

**Table S3**. Statistics for differentially methylated regions (DMRs) among the four *Aquilegia* lineages or species. Odds ratio estimates the relative methylation level between two lineages or species being compared in corresponding region. DMRs were sorted by genomic coordinates with hypo-methylated DMRs in the first lineage/species preceding hyper-methylated DMRs.

**Table S4**. Statistics for differentially methylated genes (DMGs) among the four *Aquilegia* lineages/ species. Only genes harboring a high density of differentially methylated regions (> 2 per kb) were considered DMGs.

**Table S5**. Information of the 36 *Aquilegia* samples used in this study.

