## Supplementary Notes for "Genetic and epigenetic characteristics associated with the rapid radiation of *Aquilegia* species"

1 Supplementary Notes

**Differential methylation analysis**

Differential methylation analysis was performed at the chromosome-level and the gene-level. Chromosome-level methylation similarity was measured using a chromosomal methylation discrepancy index (MDI) originally defined by O'Sullivan et al., given by

$$MDI_i = \frac{\sum_{j=1}^{N_i} |A_j - B_j|}{N_i}$$

where $\mathrm{i}$ represents the $\mathrm{i}$th chromosome containing $N_\mathrm{i}$ CG loci, while $A_\mathbf{j}$ and $B_\mathbf{j}$ represent methylation levels at $\mathrm{j}$th CG locus on chromosome $\mathrm{i}$ of two clades (lineages or species) respectively. To calculate clade-specific methylation level of one CG locus, all samples of the same clade were pooled for methylated read counts and unmethylated read counts separately. The resulting total methylated read counts and total unmethylated read counts were used to derive an integrated methylation $\beta$ value. CG loci where not both clades had ≥ 10 total read counts were excluded from MDI calculation to reduce noise. Differentially methylated genic regions (DMRs) were detected by Cochran-Mantel-Haenszel (CMH) test, which originally intends to test for weighted association of a series of $2 \times 2$ tables. CMH test handles stratified data ideally having a common odds ratio, which could be applied to tiled CG methylation levels given methylation levels at adjacent CG loci are known to be highly correlated. Specifically, when comparing two clades, samples of the same clade were first pooled with total read depth threshold at 10 as described above, and each valid CG locus provided one $2 \times 2$ table. CMH test was performed on a basis of 200bp sliding window covering all genic regions starting at the transcription start site (TSS; assumed to be the first base from the 5' end of the 5' UTR) and terminating at the transcription end site (TES; assumed to be the first base from the 3' end of the 3' UTR). Since gene-level differential methylation was of our interest, only regions containing ≥ 6 valid CG loci (three CG dinucleotides) were retained. Significantly differentially methylated genic regions were defined as having a Benjamini-Hochberg adjusted p value < 0.05 and a between-clade overall methylation level difference > 0.25 across all valid CG loci. For a differentially methylated gene (DMG), a considerable proportion of its genic sequence should be DMR. Thus, we defined DMR density as number of DMR per kilobase (kb). DMGs were required to have a DMR density ≥ 3. Since differential methylation analysis is known for false positive inflation, statistically identified DMGs were visually verified before biological interpretation using Integrative Genomics Viewer.

**Causal effect of CG mutation on epigenetic variability**

588,659 CG loci previously used for decomposition of methylomic population structure were categorized into two groups depending on whether the CG dinucleotide *per se* carried any CG-loss mutations. A linear regression model was adopted to measure the direct association between CG mutation and CG methylation loss for 224,222 mutation-carrying CG loci respectively, which was formulated as

$$M = \alpha + \gamma G + \mathrm{c}$$

where $M$ represents methylation level at a specific CG locus; $G$ represents categorical genotype of the same CG locus being 0 (homozygous non-mutant), 1 (heterozygous mutant) or 2 (homozygous mutant) with coefficient $\gamma$; $\alpha$ (intercept) represents baseline methylation level; and $\mathrm{c}$ is random error. For each CG locus, epigenetic variability explained by genetic variatio *per se* was measured by $R^2$ of the linear model.

**Identification of *cis*-driver mutations of DMRs**

For each *A. japonica - A. oxysepala* DMR, point mutations occurring inside the region or within the range of 500bp upstream/downstream were gathered from the 3,075,810 common point mutations pre-selected based on quality, missingness and minor allele frequency thresholds. All CG loci in each DMR were gathered respectively from the 588,659 high-quality CG loci for each of the 36 samples. Consequently, the selection criterion for valid CG loci was stricter than that adopted for differential methylation analysis between two clades and most DMRs did not contain the same amount of valid CG loci. To maintain sufficient DMRs in subsequent analysis, DMRs were only required to contain two valid CG loci. For each of 1,229 retained DMRs, a principal component analysis (PCA) was performed on all CG loci and the first principal component ($PC_1$) was extracted to represent the overall methylation level. An Eigenstrat method was adopted to estimate the effect of any driver mutation:

$$PC_{1_{DMR}} = \alpha + \beta_1 PC_{1_{pop}} + \beta_2 PC_{2_{pop}} + \mathrm{c}$$

where $PC_{1_{DMR}}$ stands for local methylation level-derived $PC_1$, $PC_{1_{pop}}$ and $PC_{2_{pop}}$ account for inherent population stratification which are genotype-derived $PC_1$ and $PC_2$ previously obtained from a representative LD-pruned set consisting of 15,988 common point mutations. Residual c of each model was then regressed on local genotype:

$$\mathrm{c} = \gamma G_{DMR} + \epsilon_{\mathrm{G}}$$

where $G_{DMR}$ consists of point mutations inside and around each DMR as described above. Due to limited sample size in this study, a threshold for genome-wide statistical significance was difficult to determine. Thus, differentially stringent genome-wide p value cutoffs at $5 \times 10^{-5}$, $5 \times 10^{-8}$, and $5 \times 10^{-11}$ were adopted to identify potential driver mutations.
